# Maximize crop production and environmental sustainability: insights from an ecophysiological model of plant-pest interactions and multi-criteria decision analysis

**DOI:** 10.1101/2022.02.28.482328

**Authors:** Marta Zaffaroni, Daniele Bevacqua

## Abstract

Satisfying the demand for agricultural products while also protecting the environment from negative impacts of agriculture is a major challenge for crop management. We used an ecophysiological model of plant-pest interaction and multi-criteria decision analysis to optimize crop management when considering two contrasting objectives: (1) maximizing crop production and (2) minimizing environmental impact related to fertilization, irrigation and pesticide deployment. The model provides an indicator of crop production for 27 management scenarios, obtained combining three levels of fertilization, irrigation and pesticide use, respectively. We computed the environmental impact relevant to each management scenario by means of a weighted sum of costs assigned to fertilization, irrigation and pesticide use. We identified the optimal scenarios with respect to the considered objectives analysing the Pareto front. These scenarios were mostly characterized by high fertilization and no pesticide use. We evaluated the multi-functionality of the optimal scenarios by mean of the Gini coefficient: the scenario better assuring the equality between the two objectives was characterized by high fertilization, intermediate irrigation and no pesticide. Although our results remain qualitative and not immediately transferable to agronomic practices, our analytical framework provides a useful tool to evidence trade-offs among two contrasting objectives and provide solutions to act in an efficient way by leaving a certain degree of freedom to the political decision maker.

## 1. Introduction

The world population is continuously increasing and, according to estimates, it would be 1.5 times the current population by 2100 (Li et al., 2019; Deknock et al., 2019). To ensure global food security, an increase in crop yield is essential (Li et al., 2019). Industrial forms of modern agriculture aim to remove limitations to plant productivity mainly by increasing *i*) chemical fertilizers application, to meet plants’ nutrient needs (Gliessman, 2015; Bommarco et al., 2013); *ii*) irrigation, to meet plants water needs which might increase due to warming temperature (Turner et al., 2019); *iii*) pesticide use, to control pests, which reduce crop productivity, directly by feeding and indirectly by transmitting viruses (Deknock et al., 2019; Oerke, 2006).

The way how agricultural crops are managed can significantly affect the environment and biodiversity (Bommarco et al., 2013; Seppelt et al., 2013; Demestihas et al., 2017). Mineral fertilization declines the quality of drained water, resulting in a series of environmental impacts that include surface water eutrophication, groundwater pollution, and soil degradation (Demestihas et al., 2017; Li et al., 2019). Furthermore, common fertilizer tends to volatilize into the air in the form of N_2_O, a greenhouse gas with global warming potential (GWP) 298 times that of the reference gas, CO_2_ (Xiao et al., 2019). The agricultural sector is the largest consumer of water: agriculture uses 69% of all freshwater withdrawals (UNESCO, 2020). Exploitation of both surface and ground water resources to sustain crops irrigation can reduce water flow to rivers, lakes and wetlands, and causes sustained drawdown in aquifer head levels (Calzadilla et al., 2010; Turner et al., 2019). The uneven distribution of water (and population) among world regions has made water supply critical for a growing number of countries: this trend is likely to be exacerbated by climate change (IPCC, 2021). The accumulation of pesticide residues in the environment is a threat for human and environmental health, accounting for the contamination of drinking water supplies and food sources and the reduction of biodiversity (Deknock et al., 2019). In addition, pesticide may impair bee colonies, consequently decreasing pollination, an essential service for crop production, the value of which has been estimated at €153 billion (Gallai et al., 2009).

Satisfying crop demand while guaranteeing other ecosystem services represents one of the greatest challenges facing crop management. Yet, the presence of several interacting components and biophysical processes makes the relationships between agricultural practices, crop production and environmental impact not straightforward (Demestihas et al., 2017). Models may help in attaining an understanding of agroecological systems and processes (Grimm, 1994) and they can be used to analyse the effects of a wide range of management scenarios, supplementing field experiments for identifying best management strategies (Malik and Dechmi, 2020). For example, Malik and Dechmi (2020) used a crop growth model to simulate the effect of different nitrogen management practices on maize, wheat, barley, sunflower and alfalfa fields, considering crop production and N losses. Demestihas et al. (2018) used a crop model to explore how agricultural management practices, such as planting density, irrigation and fertilization, affect the production of ecosystem services by altering soil fertility, greenhouse gas emission, water quality and fruit production in apple orchards.

Production and environmental objectives are often contrasting. One approach for evaluating management alternatives facing conflicting objectives is multi-criteria decision analysis (MCDA). It makes use of formal methods to structure and formalize the comparison of multiple objectives to support the decision making (Belton and Stewart, 2002). MCDA has been widely and successfully applied to agriculture. For example, Scharfy et al. (2017) applied MCDA to rank different clean technologies in agriculture (*e*.*g*. integrated pest management or drip irrigation) with respect to ecological, economical and social benefits. Chukalla et al. (2018) applied MCDA to evaluate the trade-off between irrigation water consumption and water pollution under different nitrogen application rates. Balezentis et al. (2020) proposed an integrated approach, combining mathematical programming and MCDA, to evaluate different scenarios to promote organic farming. MCDA has been successfully applied also to forest (Kangas and Kangas, 2005; Diaz-Balteiro and Romero, 2008; Schwenk et al., 2012; Lafond et al., 2017), and to city and tourism (Ferretti et al., 2014; Michailidou et al., 2016; Pesce et al., 2018; Langemeyer et al., 2020) management. Crucially, giving the possibility to combine economic, ecologic and social criteria, MCDA is well suited to address interdisciplinary and complex environmental questions, such those emerging in the management of agricultural systems.

In the current study, we use an ecophysiological model, presented in Zaffaroni et al. (2020), to estimate crop production and environmental costs under different management scenarios (*i*.*e*. combination of different levels of fertilization, irrigation and pesticide use). Then we apply MCDA to support the overall goal of managing agricultural crop with respect to two contrasting objectives: to maximize crop production and to minimize environmental impact. In particular, we highlight optimal alternatives respect to the two objective trough the analysis of the Pareto-front. Our work shows that the integration of a mathematical model and MCDA, giving the possibility to combine economic and ecologic objectives, is well suited to address interdisciplinary and complex environmental questions, such those emerging in the management of agricultural systems.

## 2. Materials and methods

### 2.1. Overview of the methodology

An overview of the methodology used in the present work is presented in Figure 1. We define 27 scenarios, consisting in combination of different levels of fertilization, irrigation and pesticides levels (section 2.3). The agronomic practices accounted in the different scenarios are translated in as many parameters combinations that are used: *i*) in an ecophysiological model (section 2.2), to compute the “Biomass loss” (equation 2) and *ii*) in an environmental model, to compute the “Environmental impact” (equation 3). Finally, the managements scenarios are compared via MCDA, to define the pareto optimal scenarios (section 2.5).

**Figure 1:**
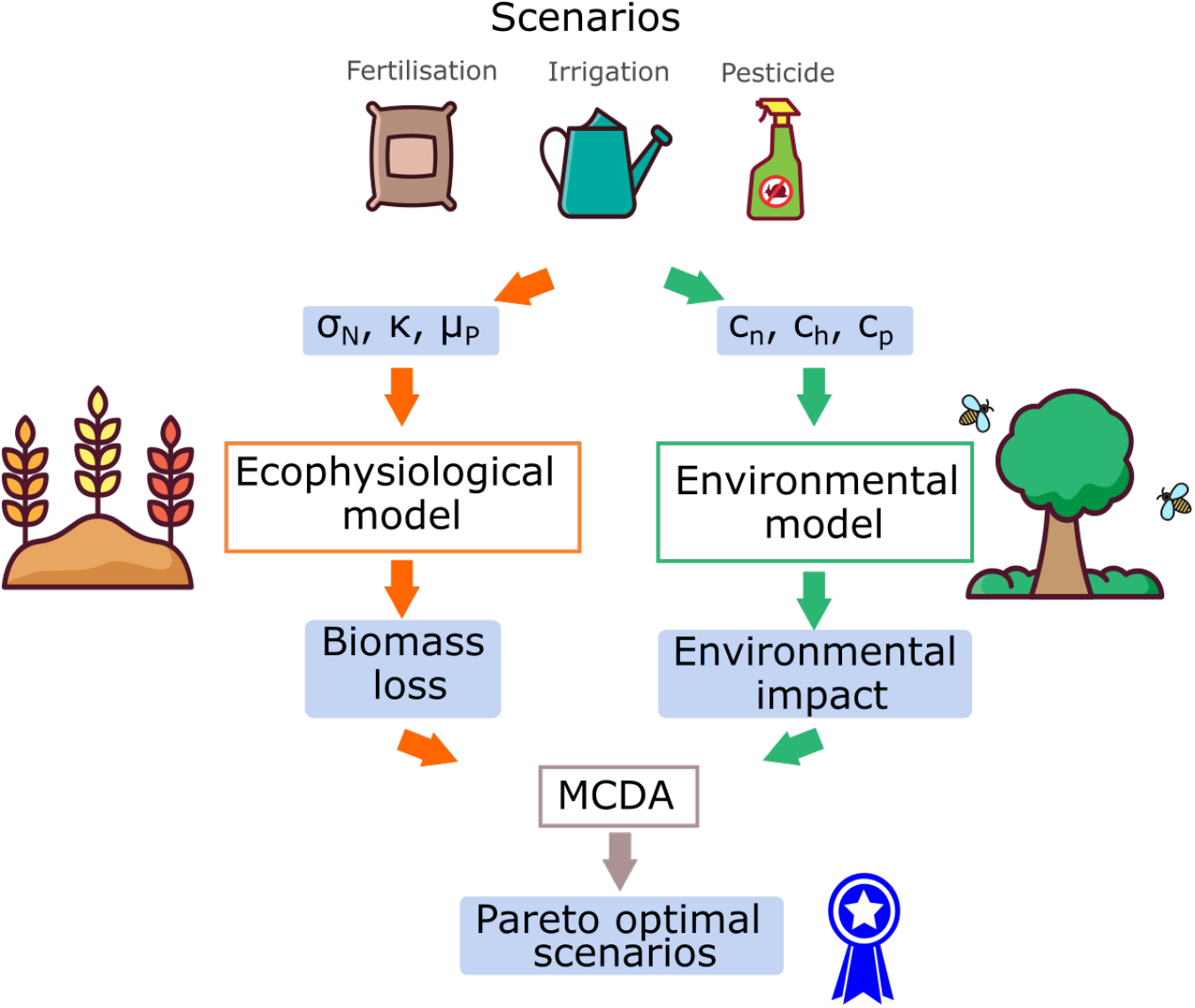
Overview of the methodology presented in the paper. An integrated approach, combining mathematical models and MCDA is used to evaluate different management scenarios with respect tot two contrasting objectives (minimizing biomass loss and environmental impact).

### 2.2. The ecophysiological model

The ecophysiological model is described in details elsewhere (see Zaffaroni et al. 2020). We provide here only an overview of its main features, schematically represented in Figure 2. The model describes the temporal variation during a growing season of *i*) dry biomass of total shoots and roots (*S* and *R*) of an average plant, *ii*) carbon and nitrogen substrates in shoots and roots (*C*_*j*_, *N*_*j*_ *j* = *S, R*), *iii*) plant induced defensive compounds (*D*), and *iv)* aphid population feeding on plant phloem and impairing plant growth (*A*). Plant is induced by aphids presence to divert carbon and nitrogen substrates from growth to defence (Will et al., 2013; Zust and Agrawal, 2016; Vyska et al., 2016). This results in the production of chemical and morphological/physiological changes that reduces aphid accessibility to the phloem (e.g. by phloem sealing) (Medina-Ortega and Walker, 2013; van Velzen and Etienne, 2015) and decreases the rate at which ingested food is converted into progeny (e.g. by releasing toxic components in the sieve that can even repel or kill the aphids) (Zust and Agrawal, 2017). The rationale behind model assumptions is discussed in detail in Zaffaroni et al. (2020). The model have been calibrated for the system peach *Prunus persica*-green aphid *Mizus persicae*. We describe the dynamics of the plant-aphid system with the following system of ordinary differential equations, the list of the considered processes (in capital Greek letters in the model) along with the list of parameters can be found in the *Supplementary Information* :

**Figure 2:**
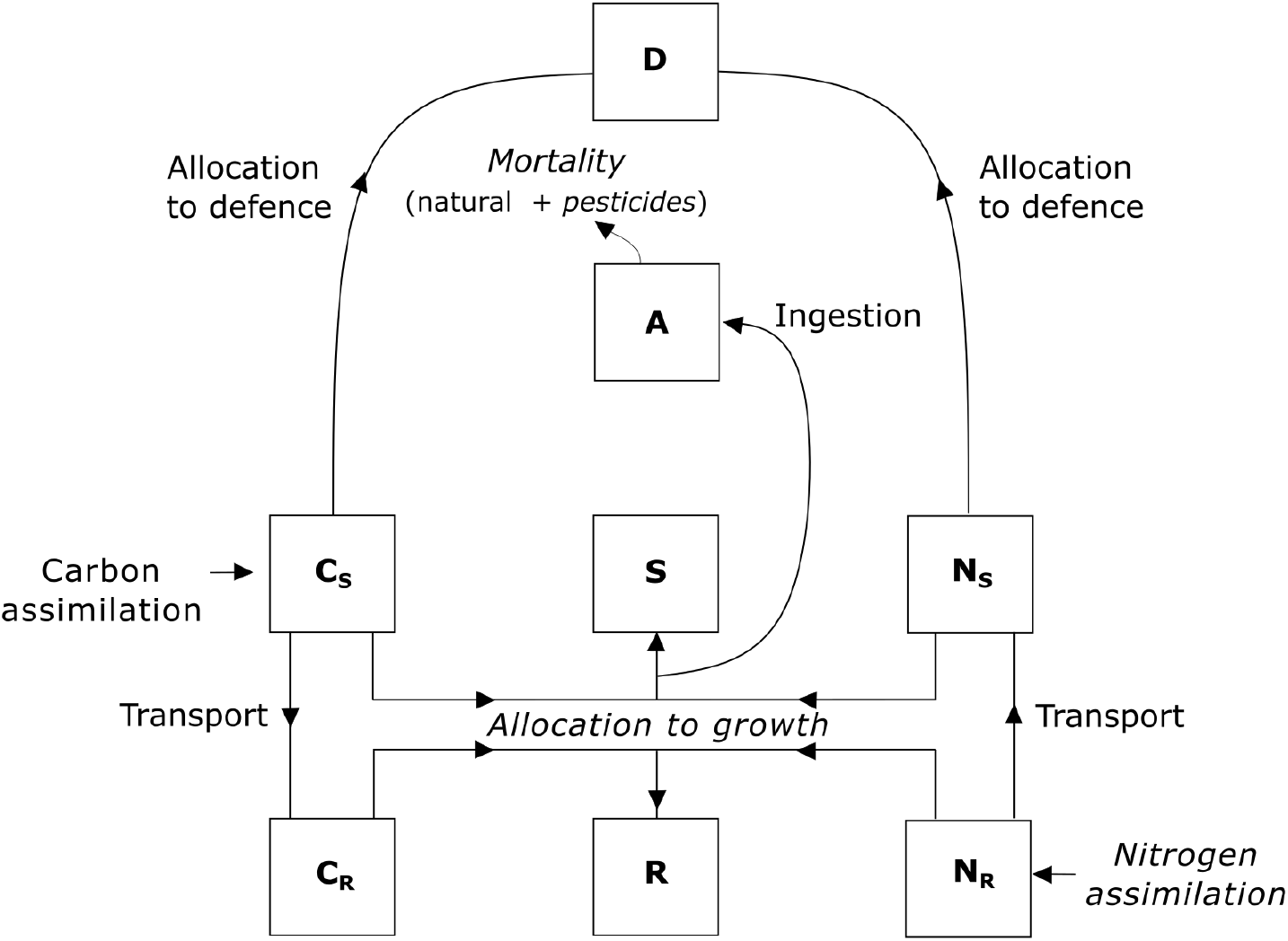
Schematic representation of the plant-aphid model where the plant is constituted by shoot (*S*) and root (*R*) structural dry mass, carbon (*C*_*j*_) and nitrogen (*N*_*j*_) substrates in shoots (*j* = *S*) and roots (*j* = *R*). The aphid population (*A*) intercepts a fraction of substrates allocated to constitute shoot structural mass and the plant diverts shoot substrates (carbon and nitrogen) to produce defences compounds (*D*). Fertilization increases nitrogen assimilation; irrigation increases plant allocation to growth, and pesticides use increases aphid mortality. More details are given in the main text.

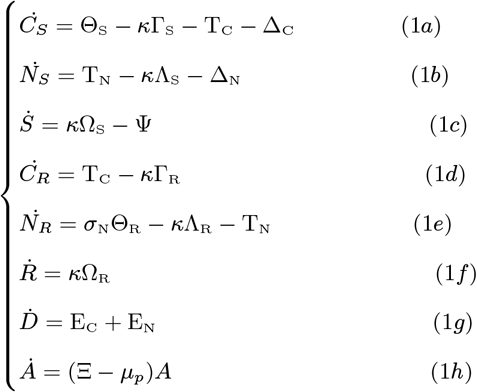

In equation (1a), Θ_S_ is the carbon assimilated by shoots from the atmosphere via photosynthesis and stored in shoots. The term *κ*Γ_S_ is the carbon substrate allocated to shoot growth or reserves. The parameter *κ* is the rate of substrate utilization for shoot biomass growth, which we assume to decrease in water stress condition (Muller et al., 2011; Sevanto, 2014). The term T_*C*_ is the carbon substrate transported from shoots to roots, which depends on carbon concentration difference between shoots and roots divided by a resistance, and Δ_*C*_ is the carbon substrate allocated to produce induced defences, which is proportional to aphid abundance. In equation (1b), T_*N*_ is the nitrogen substrate transported from roots to shoots, which, similar to carbon transport, depends on nitrogen concentration difference between roots and shoots divided by a resistance. The term *κ*Λ_S_ is the nitrogen substrate allocated to shoot growth or reserves, and Δ_*N*_ is the nitrogen substrate allocated to produce induced defences, which is proportional to aphid abundance. In equation (1c), *κ*Ω_S_ is the increase in structural shoot dry mass in the absence of any phloem withdrawal by the aphids: it depends on the rate of substrates utilization for shoots growth and it accounts for the suspension of plant growth driven changes in the photo-period. The terms Ψ represents the amount of phloem ingested by aphid population on plant, which decreases with plant defences. The dynamics of carbon and nitrogen substrate in roots (*C*_*R*_ and *N*_*R*_) and of roots dry mass (*R*) follow similar rules for nitrogen substrate assimilation, transport and allocation to root growth and we assumed that they are not directly affected by the presence of aphids. In equation (1e), the term *σ*_*N*_ Θ_*R*_ is the nitrogen assimilated by roots from the soil and stored in roots: the parameter *σ*_*N*_ is the nitrogen assimilation rate, which we assume to decrease in nutrient stress condition (Connor et al., 2011). In equation (1g), the terms E_C_ and E_N_ are the amounts of, respectively, carbon-base and nitrogen-base induced defences. In equation (1h), Ξ is the aphid intrinsic growth rate, accounting for birth rate, which increase with the ingested phloem and decrease with plant defences and with the natural mortality rate. The term *µ*_*p*_ is the pesticide induced aphid mortality rate.

We simulate the effect of variation in fertilization treatments through a variation of nitrogen assimilation rate (*σ*_*N*_), which directly affects the amount of nitrogen assimilated by roots. We simulate the effect of variation in irrigation treatment through a variation of the rate of substrates utilization for plant growth (*κ*), which directly affects the allocation of carbon and nitrogen substrates to shoot and root growth. The rationale behind these assumptions is discussed in detail in Zaffaroni et al. (2020). In the present work, we add the term for aphid pesticide induced mortality (*µ*_*p*_) which we assume to increase with pesticide.

### 2.3. Management scenarios

We consider 27 management scenarios obtained by combining three different levels (*i*.*e*. low, intermediate, high) of the three considered decision variables: fertilization, irrigation and pesticide (table 1). Note that when we refer to a low pesticide level we refer to no pesticide application. For each scenario we can simulate the trajectory of shoot dry mass *S*(*t*) and we assume that the shoot biomass at the end of the vegetative season (*S*∗ = *S*(*t* = *t*_*H*_)) is a good proxy of overall plant growth and relevant crop.

**Table 1:**
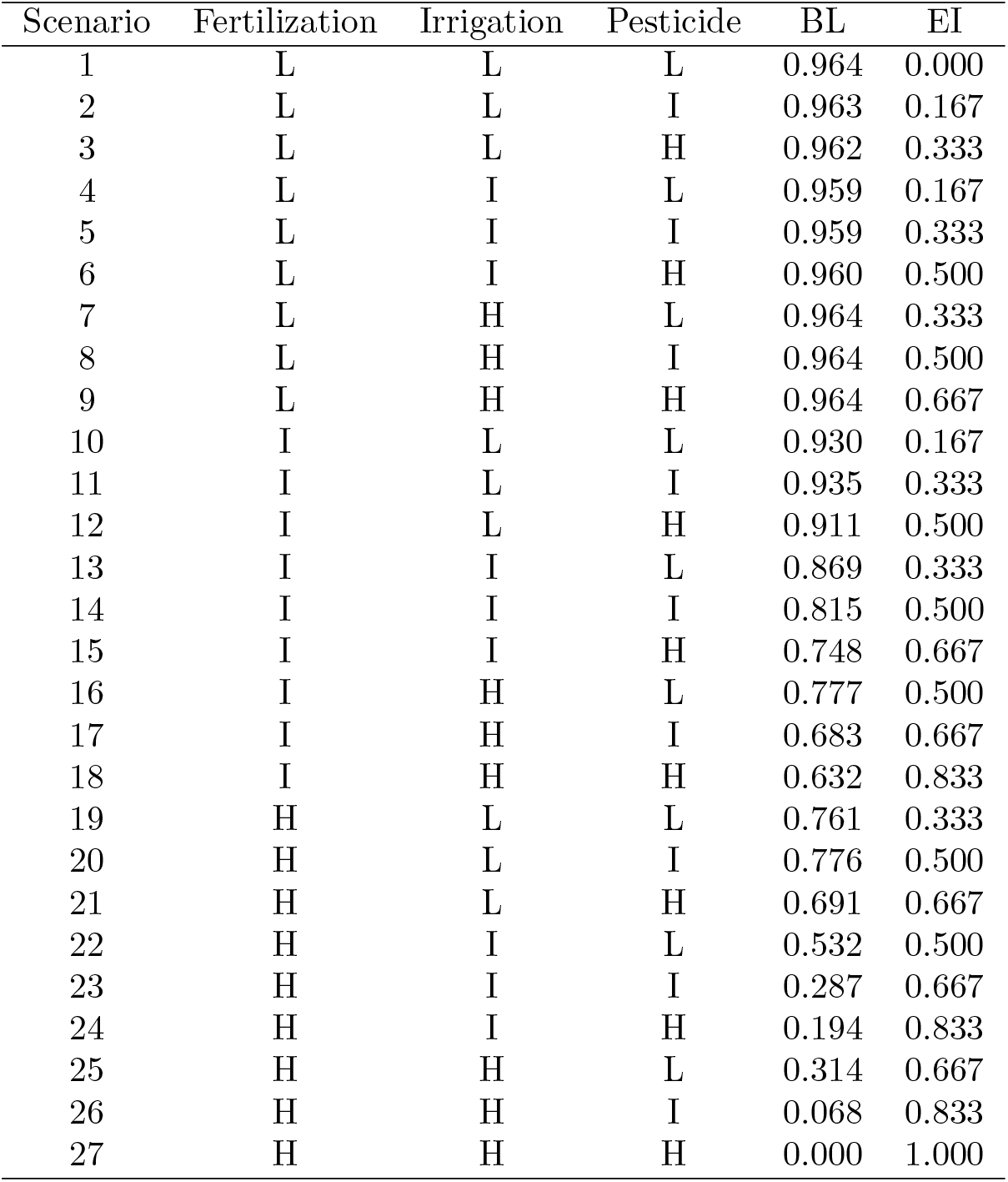
Summary of the scenarios characterized by different levels (low = L, intermediate= I, high = H) of fertilization (L: *σ*_*N*_ = 0.0012 d^-1^, I: *σ*_*N*_ = 0.012 d^-1^, H: *σ*_*N*_ = 0.12 d^-1^), irrigation (L: *κ* = 18 d^-1^, I: *κ* = 182 d^-1^, H: *κ* = 1820 d^-1^) and pesticides (L: *µ*_*p*_ = 0 d^-1^, I: *µ*_*p*_ = 0.0625 d^-1^, H: *µ*_*p*_ = 0.125 d^-1^), of the estimated biomass loss (BL) and environmental impact (EI).

### 2.4. Identification of objectives

We evaluate the performance of each management scenario with respect to two objectives: to maximize the plant biomass production and to minimize the environmental impact (EI). Since maximizing plant biomass production *S*∗ is equivalent to minimizing plant biomass loss (BL), in order to have two objectives to be minimized, we will consider the latter as objective in our analysis. We compute the plant biomass loss for the scenario *i* as the fraction of shoot biomass at the end of the vegetative season (*S*∗) lost respect to the maximum potential value:

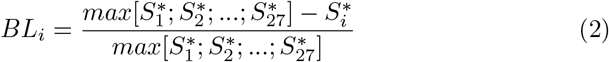

This implies that BL is equal 0 when the shoot production is maximum and is equal to 1 when the shoot production is null. For each management scenario we assign a cost (*c*_*k*_) to the three decision variables [*i*.*e*. fertilization (*k* = *n*), irrigation (*k* = *h*), pesticide (*k* = *p*)] as follows: *i*) *c*_*k*_ = 0 when human intervention is low; *ii*) *c*_*k*_ = 0.5 when it is intermediate; *iii*) *c*_*k*_ = 1 when it is high. Then, we model the environmental impact of the of the *i*^*th*^ management scenarios by mean of a weighted sum:

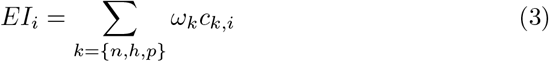

In the case that the same environmental cost is associated to fertilization, irrigation and pesticides one should give equal weights to decision variables associated to the three interventions (*i*.*e. ω*_*n*_ = *ω*_*h*_ = *ω*_*p*_ = 0.33). On the other hand, if one were more concerned with respect to one environmental cost, the weights values can be accordingly varied. As for BL, EI values range between 0 and 1, and the three weights must sum one.

### 2.5. Multi-criteria decision analysis

MCDA is a multi-step process comprising a family of methods to structure and formalize the comparison of management scenarios to support the decision-making (Pesce et al., 2018; Zanchi and Brady, 2019). We firstly explore the possible trade-off between the two objectives (minimizing BL and EI) through the identification of the Pareto front. The Pareto front specifies the groups of Pareto-optimal scenarios, *i*.*e*. the management scenarios for which it is not possible to modify the decision variables (*i*.*e*. fertilization, irrigation and/or pesticide) to improve the performance respect to one objective (i.e. decreasing BL) without worsening at the same time the performance respect to the other objective (*i*.*e*. increasing EI) (Kennedy et al., 2008).

Since the weights assigned to the decision variables influences the value of the objective EI and consequently the Pareto-optimal scenarios, we conduce a sensitivity analysis to test the robustness of the Pareto-optimal scenarios. We consider other weights combination to mimic different decision makers’ concerns relate to fertilization, irrigation and pesticide environmental impacts. We assign a cost *c*_*k*_ = 0.9 to the decision variable whose environmental impact concerns the decision makers the most, and a cost *c*_*k*_ = 0.05 to the other two decision variables. Thus, we analyze the variation in the scenarios composing the Pareto front with respect to the original MCDA evaluation, which considers equal weights for the three decision variables (“equal concern” weight combination).

For each Pareto-optimal scenario we compute the Gini coefficient, which describes the inequality in the objectives values and indicate the multi-functionality of a scenario. Although Gini coefficient has originally been employed in economic analysis as a valid index to measure the income inequalities, it has been used in ecological application as well (Accatino et al., 2018; Wang et al., 2019; Li et al., 2017). According to Dorfman (1979), we computed the Gini coefficient for each scenario *i* as :

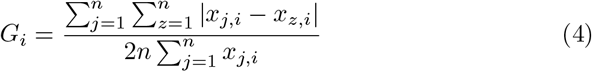

where *n* (= 2) is the number of considered objectives and *x*_*j*_ and *x*_*z*_ are the objective values for the considered scenario. A Gini coefficient of zero expresses perfect equality between the objectives, increasing the value of the Gini coefficient the inequality between the objectives increases and the multi-functionality of a scenario decreases.

## 3. Results

We report the values of the two objectives evaluated for different management scenarios in table 1 and in figure 3. Biomass loss varies between 0 and 0.96, while the environmental impact varies between 0 and 1. The Pareto front highlights that as biomass loss decreases, ecological impact increases and vice versa. A trade-off between the considered objectives clearly exists. The Pareto front is composed by 7 out of 27 considered management scenarios (hereafter the Pareto-optimal scenarios). Among these 7: the 72% is characterized by an high level of fertilization (scenarios 19, 22, 23, 26, 27), the 14% by, respectively, a low and intermediate levels of fertilization (scenarios 1 and 10, respectively); the 42.6% is characterized by a low irrigation (scenarios 1, 10, 19), the 28.7% by, respectively, an intermediate and high level of irrigation (scenarios 22, 23 and 26, 27 respectively); the 57% is characterized by a low pesticide level (scenarios 1, 10, 19, 22), the 29% by an intermediate level of pesticide (scenarios 23 and 26) and the 14% by an high level of pesticide (scenario 27).

**Figure 3:**
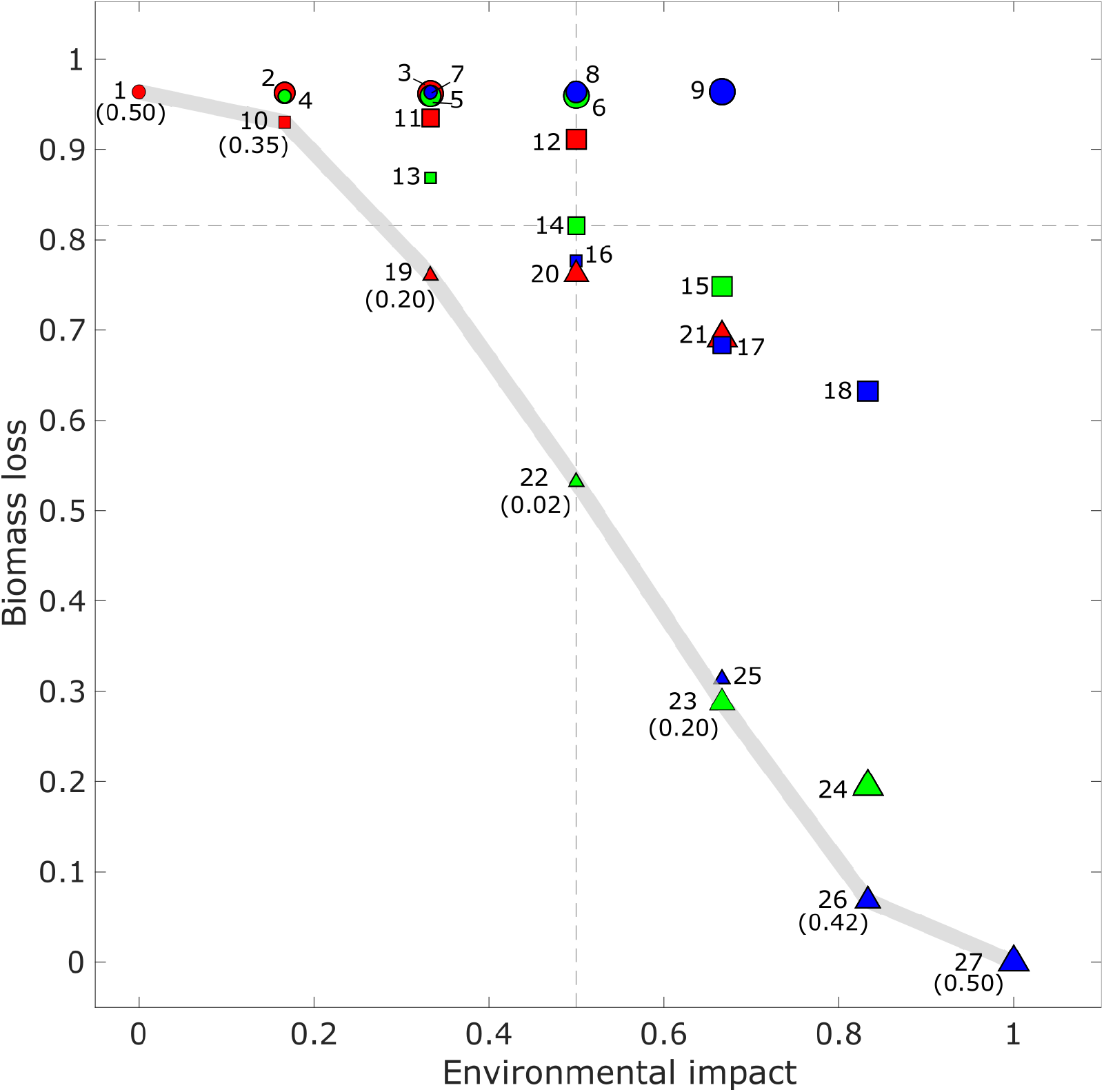
Biomass loss and environmental impact for different management scenarios characterized by different values of fertilization (parameter *σ*_*N*_) (symbol shape: circle = low, square = intermediate, triangle = high); irrigation (parameter *κ*) (symbol color: red = low, green = intermediate, blue = high); and pesticides (parameter *µ*_*p*_) (simbol size: small = low, intermediate = intermediate, big = high). The grey line represents the Pareto front. Scenario 14 is the baseline scenario characterized by intermediate levels of fertilization, irrigation and pesticide use. Dashed lines identify four group scenarios: win_P_ - win_E_ (bottom-left), win_P_ - lose_E_ (bottom-right), lose_P_ - lose_E_ (top-right), lose_P_ - win_E_ (top-left). The Gini coefficient, indicating inequality, for the optimal scenarios is indicated between parentheses.

As expected, the weights assigned to the decision variables influence the EI and consequently which scenarios are labelled as optimal. However, the sensitivity analysis showed that the scenarios composing the Pareto front obtained with the “equal concern” weight combination are optimal independently from the weights assigned to the decision variables costs (*ω*_*k*_) (see table 2). With the other weight combinations, the number of optimal scenarios increased with respect to the “equal concern” case. Yet, sensitivity analysis showed that the majority of Pareto-optimal scenarios are characterized by high fertilization (figure 1 A-G-J in *Supplementary Information*) and low pesticide (figure 1 C-F-I-L in *Supplementary Information*), independently by the weights combination. Only when the cost associated to fertilization is predominant (figure 1 D in *Supplementary Information*) the majority of Pareto-optimal scenario are characterized by intermediate fertilization level.

**Table 2:**
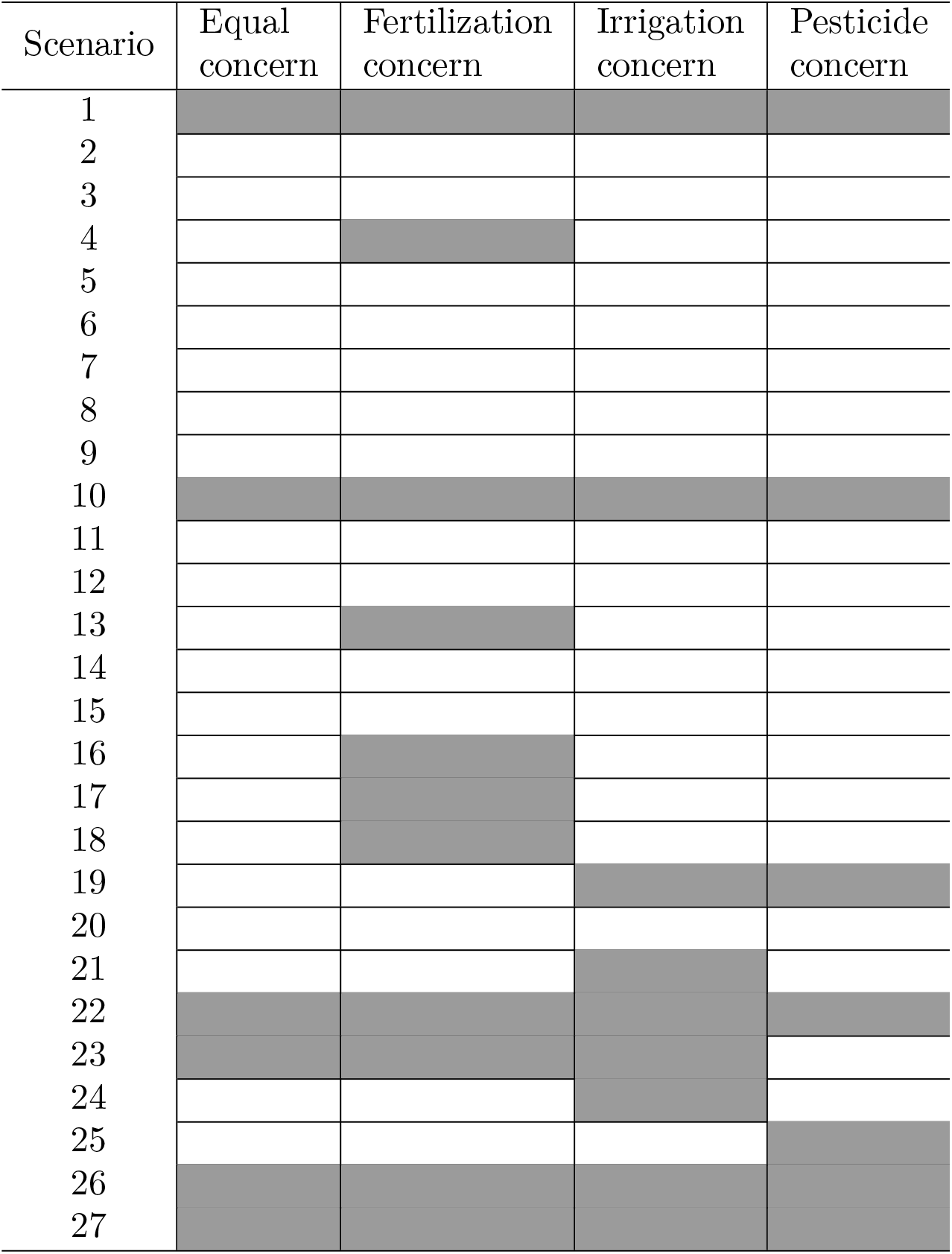
Pareto optimal scenarios (in grey) across different weights combinations: *i*) equal concern (*ω*_*n*_ = *ω*_*h*_ = *ω*_*p*_ = 0.33); *ii*) fertilization concern (*ω*_*n*_ = 0.9, *ω*_*h*_ = *ω*_*p*_ = 0.05); *iii*) irrigation concern (*ω*_*h*_ = 0.9, *ω*_*n*_ = *ω*_*p*_ = 0.05); *iv*) fertilization concern (*ω*_*p*_ = 0.9, *ω*_*n*_ = *ω*_*h*_ = 0.05).

The baseline scenario, which is characterized by an intermediate value for fertilization, irrigation and pesticides application (scenario 14 in figure 3), is not an optimal scenario. Starting from the baseline scenario, we identify four scenarios groups: *i*) win_P_ - win_E_ scenarios, where both the (P)roduction and the (E)nvironmental objectives are improved (or do not change) respect to the baseline scenario; *ii*) win_P_ - lose_E_ scenarios, where BL decreases (or does not change), but EI increases; *iii*) lose_P_ - win_E_ scenarios, where EI decreases (or does not change), but BL increases; *iv)* lose_P_ - lose_E_ scenarios, where both the objectives are worsened respect to the baseline scenario. The win_P_ - win_E_ scenarios (*i*.*e*. scenarios 16, 19, 20, 22) are the most interesting and, in our case, they are all characterized by an higher fertilization level with respect to the baseline scenarios.

The values of Gini coefficient for the Pareto-optimal scenarios are reported between parenthesis in figure 3. Scenario 22 is the one which better assures the equality between the environmental and the production objectives. It is characterized by high fertilization, intermediate irrigation and low pesticide, as expected. Scenarios 1 and 27 presents the highest inequality between the objectives: in scenario 1 the environmental objective dominates, while in scenario 27 the production objective dominates.

## 4. Discussion

In this work, we show that a mechanistic model reproducing plant-aphid interaction can be used to simulate the impact of agricultural practices on a plant-aphid system. The strengths of our model are, arguably, that it is relatively simple and transparent, providing clear connections between assumed mechanisms and predicted response. Transparency is important, so that complex and sometimes unexpected interactions between agricultural practices, plant growth and aphid population dynamics can be disentangled. For example, our model assumes that fertilization has a double role: it fosters both plant growth (Robertson and Vitousek, 2009) and the production of induced defences, which eventually reduces aphid pressure on the plant (Zust and Agrawal, 2016). This mechanism likely explains why the scenarios with the highest predicted yield are obtained with high fertilization. Moreover, our model assumes that both nitrogen and carbon are essential to plant to produce new biomass. This likely explains why, increasing 10 times nitrogen assimilation rate, without varying carbon assimilation rate depending on water availabilty (parameter *k*), does not result in a proportionate increase in the production of new plant biomass: for example by increasing of 10 times (*i*.*e*. scenario 18) the fertilization level we obtained an increase in biomass production of 2.7 times. However, our results should be interpreted as qualitative pattern and they cannot be immediately transferred to agronomic practices. In fact, to derive effective tools for decision analysis, one should define the functional relationships between model parameters and actual agronomic practices. Models that linked soil nitrate content to nitrogen uptake by crop have been proposed (see e.g. Cárdenas-Navarro et al. 1999; Devienne-Barret et al. 2000): further works should investigate how this type of models can be adapted to be used in mechanistic plant models to reproduce different fertilization treatments.

Our work shows a trade-off between plant biomass loss and environmental impact of the considered management scenarios. Such trade-off is a common finding in both empirical (Jiang et al., 2013; Raudsepp-Hearne et al., 2010;Tilman et al., 2002) and simulation (Goldstein et al., 2012; Nelson et al., 2009; Kirchner et al., 2015) works. Pareto-optimal scenarios are mostly characterized by high fertilization level. The environmental impacts linked to fertilization are mainly due to the fact that crops take up only about 30%-50% of nitrogen fertilization applied leaving most of the remainder available for loss by volatilization and leaching (Tilman et al., 2002; Malik and Dechmi, 2020). Agricultural practices that allow to increase nitrogen use efficiency, *i*.*e*. the fraction of applied nitrogen that is absorbed and used by the plant (in our model this would translate into higher vales of *σ*_*N*_), represents a promising solution to increase plant biomass production and reduce the environmental impact of fertilization (Tilman et al., 2002; Malik and Dechmi, 2020). For example, fertilize applications during periods of greatest crop demand, in close proximity of plant roots, and in smaller and more frequent doses have the potential to improve nitrogen use efficiency, reducing fertilizer losses while maintaining or improving yields and quality (Tilman et al., 2002; Cui et al., 2008; Xiao et al., 2019).

The majority of Pareto optimal scenarios is characterized by no pesticide addition. This is likely due to the fact that aphid’s feeding effect on plant biomass production is modest respect to those of other pest (e.g. defoliators) (Van Emden and Harrington, 2007). Yet, optimal scenarios considering no pesticide are characterized by medium to high plant biomass loss, thus producers may be reluctant to implement this prescription fearing for economical losses. Nevertheless, as the willingness to pay of the consumers might increase for products issued from sustainable agronomic practices that minimize the use of pesticide, differences in economical return per unit of product between the organic alternative and conventional farming may be small (Pimentel and Burgess, 2005). In the event that producers are more oriented towards scenarios minimizing biomass loss, pesticide control is necessary. In this case, agricultural practices such as biological control have the potential to limit pest outbreaks acting on aphids mortality (*µ*_*p*_) through predation or parasitism (Murdoch et al., 1985), without having the environmental impact due to pesticide.

Regular deficit irrigation (RDI) is a water-saving practice, which has been widely applied in field crops (e.g. wheat, maize, cotton, rice, soybean, sunflower) and woody plant species (e.g. grapevine, nectarine, olive trees, apple) with the aim of reducing water application without decreasing the yield (see Chai et al. 2016 and citations therein). This kind of practice allows to increase plant biomass production and reduce the environmental impact due to irrigation. Moreover RDI has the potential to control pest pressure (see e.g. Costello 2008; Mercier et al. 2008; Rousselin et al. 2016; Bevacqua et al. 2019) which, if accompanied by a lower pesticide application, may further increase plant biomass production and reduce environmental impact. Yet, timing and the extent to which RDI is applied plays a critical role in determining yield production and pest control (Chai et al., 2016).

While the methodology we present is flexible enough to apply in many contexts, we should point out some several considerations in interpreting the results of this analysis. First, we use a plant-pest model general enough to be applied to different plant-pest system. We calibrate it for young peach tree without fruits to compute the plant biomass loss objective (see Zaffaroni et al. 2020). The assumption we make to use shoot production as a proxy for relevant crop is generally true for annual plants, while for perennial plants is true if we consider the average production in the whole lifespan of plants, because they requiring a number of growth cycles before fruit is produced. Another consideration involves the assumption of independent effects of fertilization and irrigation on plant growth: by contrast, in drought stress condition N uptake by root may be restricted (McDonald and Davies, 1996). A water model (e.g. that presented in Thornley (1996)), describing the water movement from the soil to the shoot can *i*) be incorporated to our plant-pest model, to explicitly take into account the interaction between nitrogen uptake and soil water content, or *ii*) provide suggestions for empirical relationships between parameters.

## 5. Conclusion

Despite the simplifying assumptions outlined above, our work offers new insights for sustainable crop management, demonstrating the utility of analytical approaches that combine plant simulation modelling with MCDA. We do not expect that the qualitative findings of our work can immediately be transferable to agronomic practices, yet the analytical framework introduced here can be readily modified to consider different plant-pest system, to include others agricultural practices and to incorporate alternative plant models, providing a basis for further study.

