## Supplementary Information for "Maximize crop production and environmental sustainability: insights from an ecophysiological model of plant-pest interactions and multi-criteria decision analysis"

### S1 Model processes and parameters

The processes in equation 1 in the main text are reported below. Details of state variables and parameters are reported in Table S1.

$$\Theta_S = \sigma_C S \left[ \left(1 + \frac{S}{\nu}\right)\left(1 + \frac{C_S}{S\iota_{\mathrm{C}}}\right) \right]^{-1} \tag{S1}$$

$$\Gamma_S = \varphi_C \frac{C_S}{S}\frac{N_S}{S} S \tag{S2}$$

$$\mathrm{T}_C = \left(\frac{C_S}{S} - \frac{C_R}{R}\right)(SR)^q \cdot (S^q + R^q)^{-1} \tag{S3}$$

$$\Delta_C = \alpha \frac{C_S}{S} A \tag{S4}$$

$$\mathrm{T}_N = \left(\frac{N_R}{R} - \frac{N_S}{S}\right)(SR)^q \cdot (S^q + R^q)^{-1} \tag{S5}$$

$$\Lambda_S = \varphi_N \frac{C_S}{S}\frac{N_S}{S} S \tag{S6}$$

$$\Delta_N = \alpha \frac{N_S}{S} A \tag{S7}$$

$$\Omega_S = \frac{\lambda^\eta}{\lambda^\eta + t^\eta}\frac{C_S}{S}\frac{N_S}{S} S \tag{S8}$$

$$\Psi = \begin{cases} \theta A & \text{if } \theta \cdot A \leq \kappa \Omega_S \left(1 - \beta_1 \frac{\frac{D}{S}^{\delta_1}}{\pi_1^{\delta_1} + \frac{D}{S}^{\delta_1}}\right) \\ \kappa \Omega_S \left(1 - \beta_1 \frac{\frac{D}{S}^{\delta_1}}{\pi_1^{\delta_1} + \frac{D}{S}^{\delta_1}}\right) & \text{otherwise} \end{cases} \tag{S9}$$

$$\Gamma_R = \varphi_C \frac{C_R}{R}\frac{N_R}{R} R \tag{S10}$$

$$\Theta_R = R \left[ \left(1 + \frac{R}{\nu}\right)\left(1 + \frac{N_R}{R\iota_{\mathrm{N}}}\right) \right]^{-1} \tag{S11}$$

$$\Lambda_R = \varphi_N \frac{C_R}{R}\frac{N_R}{R} R \tag{S12}$$

$$\Omega_R = \frac{\lambda^\eta}{\lambda^\eta + t^\eta} \frac{C_R}{R} \frac{N_R}{R} R \tag{S13}$$

$$\mathrm{E}_C = \varepsilon_C \alpha \frac{C_S}{S} A \tag{S14}$$

$$\mathrm{E}_N = \varepsilon_N \alpha \frac{N_S}{S} A \tag{S15}$$

$$\Xi = \begin{cases} \xi\theta\left(1 - \beta_2 \frac{\frac{D}{S}^{\delta_2}}{{\pi_2}^{\delta_2} + \frac{D}{S}^{\delta_2}}\right) - \mu & \text{if } \theta \cdot A \leq \kappa\Omega_S\left(1 - \beta_1 \frac{\frac{D}{S}^{\delta_1}}{{\pi_1}^{\delta_1} + \frac{D}{S}^{\delta_1}}\right) \\ \xi\kappa\Omega_S\left(1 - \beta_1 \frac{\frac{D}{S}^{\delta_1}}{{\pi_1}^{\delta_1} + \frac{D}{S}^{\delta_1}}\right)\frac{1}{A}\left(1 - \beta_2 \frac{\frac{D}{S}^{\delta_2}}{{\pi_2}^{\delta_2} + \frac{D}{S}^{\delta_2}}\right) - \mu & \text{otherwise} \end{cases} \tag{S16}$$

Table S1: Model variables and parameters.

| Variable | | Dim. | Description | |
|---|---|---|---|---|
| S | | g | Shoot structural dry mass | |
| R | | g | Root structural dry mass | |
| $C_S$ | | g | Shoot carbon substrate | |
| $C_R$ | | g | Root carbon substrate | |
| $N_S$ | | g | Shoot nitrogen substrate | |
| $N_R$ | | g | Root nitrogen substrate | |
| $D$ | | DU | Plant induced defences | |
| $A$ | | ind. | Aphid population | |
| **Parameter** | **Value** | **Dim.** | **Description** | **Source** |
| C and N assimilation | | | | |
| $\sigma_{\mathrm{C}}$ | 0.1 | $d^{-1}$ | Assimilation rate of C | Thornley (1998) |
| $\sigma_{\mathrm{N}}$ | [1.2; 12; 120] $10^{-3}$ [a] | $d^{-1}$ | Assimilation rate of N | Zaffaroni et al. (2020) |
| $\nu$ | 1000 | g | Shoot (root) mass halving substrate assimilation due to self shading (competition) | Thornley (1998) |
| $\iota_{\mathrm{C}}$ | 0.1 | / | Semi-saturation C concentration | Thornley (1998) |
| $\iota_{\mathrm{N}}$ | 0.01 | / | Semi-saturation N concentration | Thornley (1998) |
| C and N substrates allocation to plant growth | | | | |
| $\varphi_{\mathrm{C}}$ | 0.50 | / | Unit of substrate C per unit of structural dry mass | Thornley (1998) |
| $\varphi_{\mathrm{N}}$ | 2.50 $10^{-2}$ | / | Unit of substrate N per unit of structural dry mass | Thornley (1998) |
| $\kappa$ | [18.2; 182; 1820] [b] | $d^{-1}$ | Maximum rate of substrate utilization | Zaffaroni et al. (2020) |
| $\eta$ | 73 | / | Switch-off function of plant growth: steepness | Zaffaroni et al. (2020) |
| $\lambda$ | 169 | d | Switch-off function of plant growth: date of equal partitioning between growth and reserves | Zaffaroni et al. (2020) |
| Transport | | | | |
| $q$ | 0.86 | / | Plant architecture scaling parameter | Zaffaroni et al. (2020) |
| Defences development | | | | |
| $\alpha$ | 0.02 | g $d^{-1}$ $ind^{-1}$ | Allocation of substrates to defences per unit of aphid | Zaffaroni et al. (2020) |
| $\varepsilon_{\mathrm{C}}$ | 5 $10^{-2}$ | DU $g^{-1}$ | Conversion efficiency of C substrate in defences | Schoonhoven et al. (2005) |
| $\varepsilon_{\mathrm{N}}$ | 1 | DU $g^{-1}$ | Conversion efficiency of N substrate in defences | Schoonhoven et al. (2005) |
| Aphid | | | | |
| $\theta$ | 1.12 $10^{-3}$ | g $d^{-1}$ $ind^{-1}$ | Maximum food intake per aphid | Day and Irzykiewicz (1953) |
| $\xi$ | 171 | ind $g^{-1}$ | Maximum conversion efficiency of ingested food into descendants | Saguez et al. (2005) |
| $\mu$ | 0.04 | $d^{-1}$ | Aphid natural mortality rate | Gange et al. (1999) |
| $\mu_P$ | [0; 0.0625; 0.125] [c] | $d^{-1}$ | Aphid pesticide induced mortality rate | Gange et al. (1999) !!!!!!!!!!!!!!!! |
| $\pi_1$ | 8.52 $10^{-3}$ | DU $g^{-1}$ | Swich-on function of defences protected phloem fraction: defences concentration at which defences effect is half-saturated | Zaffaroni et al. (2020) |
| $\delta_1$ | 0.65 | / | Swich-on function of defences protected phloem fraction: steepness | Zaffaroni et al. (2020) |
| $\pi_2$ | 0.05 | DU $g^{-1}$ | Swich-on function of defences induced mortality/repulsion rate: defences concentration at which defences effect is half-saturated | Zaffaroni et al. (2020) |
| $\delta_2$ | 118 | / | Swich-on function of defences induced mortality/repulsion rate: steepness | Zaffaroni et al. (2020) |
| $\beta_1$ | 1 | / | Swich-on function of defences protected phloem fraction: asymptotic value | Zaffaroni et al. (2020) |
| $\beta_2$ | 28 | / | Swich-on function of defences induced mortality/repulsion rate: asymptotic value | Zaffaroni et al. (2020) |

[a] Values refer to low, intermediate and high fertilization treatments, respectively.
[b] Values refer to low, intermediate and high irrigation treatments, respectively
[c] Values refer to no, intermediate and high pesticide treatments, respectively

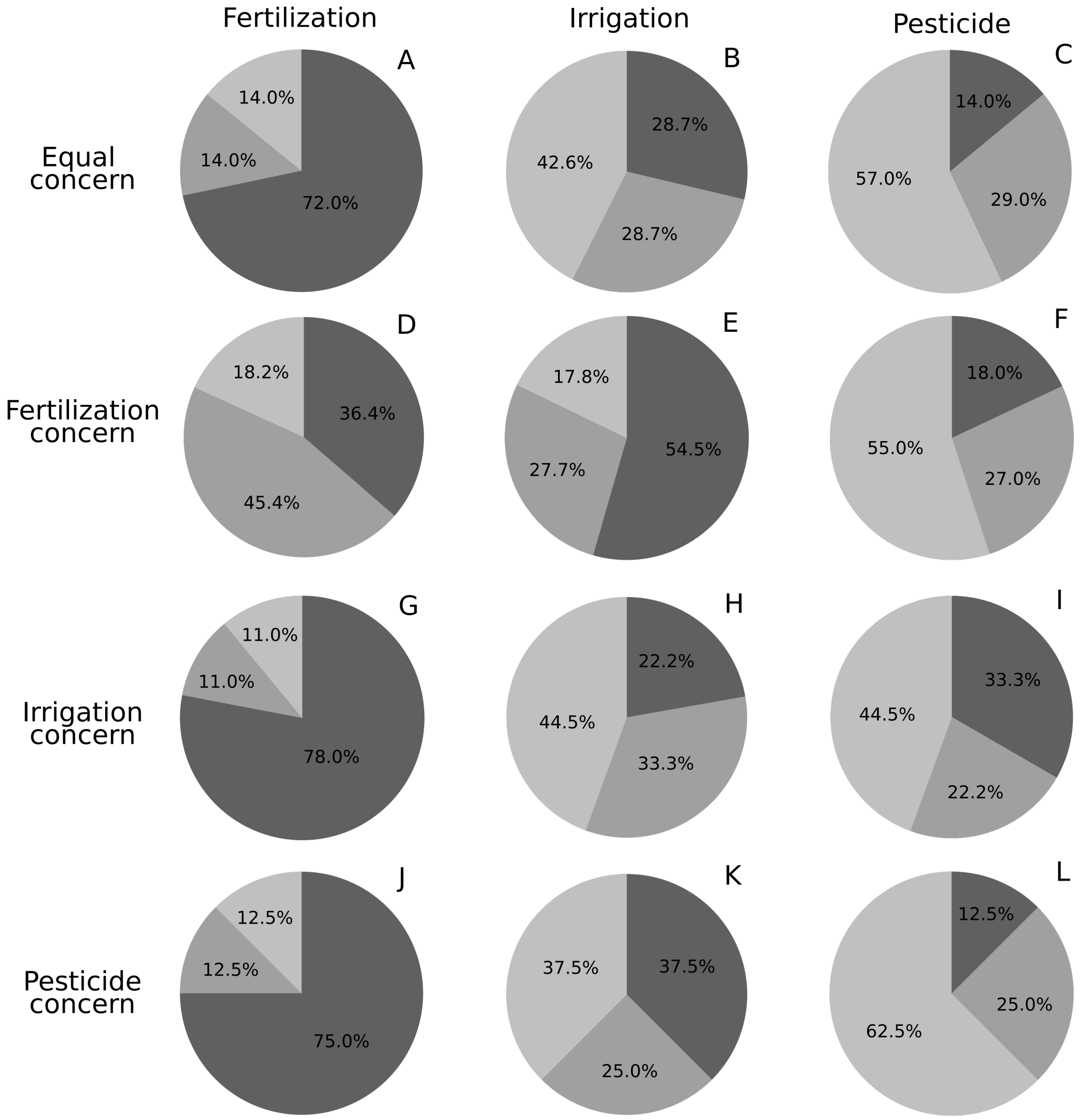


Figure S1: Response of the percentage of Pareto-optimal scenarios characterized by low (light grey), intermediate (grey) and high (dark grey) fertilization, irrigation or pesticide applications to weights combinations ($w_n, w_h$ and $w_p$). In the equal concern case (A-B-C) $w_n = w_h = w_p = 0.33$, in the fertilization concern case (D-E-F) $w_n = 0.9$, $w_h = w_p = 0.05$, in the irrigation concern case (G-H-I) $w_h = 0.9$, $w_n = w_p = 0.05$, and in the pesticide concern case (J-K-L) $w_p = 0.9$, $w_n = w_h = 0.05$.
